# IFN-induced protein with tetratricopeptide repeats 2 (Ifit2) limits autoimmune inflammation by regulating myeloid cell activation and metabolic activity

**DOI:** 10.1101/2022.03.11.483954

**Authors:** Dongkyun Kim, Nagendra Kumar Rai, Amy Burrows, Sohee Kim, Ajai Tripathi, Samuel E. Weinberg, Ranjan Dutta, Ganes C. Sen, Booki Min

**Author notes:** Corresponding author: Booki Min, Department of Microbiology and Immunology, Northwestern University Feinberg School of Medicine, 310 E Superior St. Morton 6-626. Chicago, IL 60611. EMAIL).

## Abstract

Besides anti-viral functions, Type I IFN expresses potent anti-inflammatory properties and is being widely used to treat certain autoimmune conditions, such as multiple sclerosis (MS). In murine model of MS, experimental autoimmune encephalomyelitis (EAE), administration of IFNβ effectively attenuates the disease development. However, the precise mechanisms underlying the treatment remain elusive. In this study, we report that IFN-induced protein with tetratricopeptide repeats 2 (Ifit2), a type I and type III IFN-stimulated gene, plays a previously unrecognized immune regulatory role during autoimmune neuroinflammation. Mice deficient in Ifit2 display greater susceptibility to EAE and escalated immune cell infiltration in the central nervous system. Ifit2 deficiency is also associated with microglial activation and increased myeloid cell infiltration. Unexpectedly, myelin debris clearance and the subsequent remyelination is impaired in *Ifit2*^-/-^ CNS tissues. Clearing myelin debris is an important property of reparative M2 type myeloid cells to promote remyelination. Indeed, we observed that bone marrow derived macrophages, CNS infiltrating myeloid cells, and microglia from *Ifit2*^-/-^ mice express cytokine and metabolic genes associated with proinflammatory M1 type subsets. Taken together, our findings uncover a novel regulatory function of Ifit2 in autoimmune inflammation in part by modulating myeloid cell function and metabolic activity.

## Introduction

Type I IFN (IFN-I) is widely used to treat acute and chronic inflammation including autoimmunity in the central nervous system (CNS), especially multiple sclerosis (MS) (1-4). Despite its broad applications, the precise mechanisms by which IFN-I elicits its treatment effects within the CNS remain largely unclear. In mouse model of MS, experimental autoimmune encephalomyelitis (EAE), IFNβ administration effectively ameliorates the severity of the disease (5, 6). IFN-I is also induced within the CNS during EAE, and it was reported that endogenously produced IFN-I plays a key role in regulating EAE pathogenesis, since mice deficient in IFN-I receptor (IFNAR) or in TIR domain-containing adaptor-inducing interferon-β (TRIF), a cytosolic adaptor molecule needed to induce IFN-I production, develop exacerbated EAE (7, 8). The use of cell type specific *Ifnar*^-/-^ mice further uncovers that IFN-I signaling in T cells or neurons is dispensable and that the signaling in myeloid cells was essential for the anti-inflammatory properties of endogenously produced IFN-I (7). Nevertheless, the cellular and molecular mechanisms by which endogenously produced IFN-I mediates its anti-inflammatory functions remain unknown.

IFN-I orchestrates both innate and adaptive immunity by inducing IFN stimulated genes (ISGs) in multiple cell types (9, 10). IFN-induced tetratricopeptide repeats (Ifit) is a group of ISG protein family consisted of 4 families in human (3 families in mice, Ifit1/ISG56, Ifit2/ISG54, and Ifit3/ISG60). Ifit proteins play important anti-viral roles through their ability to bind viral RNA and to interfere with translation (11-13). The anti-viral functions of Ifit overlap amongst the Ifit family proteins, although the mechanisms utilized by each Ifit member could be unique and distinct (13-15). We recently reported that Ifit2 protects mice from lethal neurotropic MHV infection (16). Pronounced mortality of *Ifit2*^-/-^ mice during the infection is accompanied by uncontrolled virus replication and spread throughout the brain (16). Interestingly, there is significantly impaired microglial activation and reduced recruitment of inflammatory cells to the brain, suggesting an Ifit2’s ability to control microglial activation. In addition to broad anti-viral functions, Ifit is known to play a diverse regulatory role in immunity. In LPS activated macrophages, Ifit1 not only supports IFN-I expression but also limits the inflammatory gene expression (17). Ifit3 negatively regulates TLR3 induced IFNβ and Cxcl10 expression in human brain endothelial cells (18). However, whether Ifit plays a role in autoimmune inflammation has not previously been examined.

In this study, we report that Ifit2 plays an important role in limiting autoimmune neuroinflammation. Mice deficient in Ifit2 were highly susceptible to EAE, and inflammatory responses in the CNS tissue were markedly elevated in these mice. Ifit2 deficiency was also associated with pronounced microglial activation and macrophage infiltration. Interestingly, severely impaired myelin debris clearance and remyelination were observed in *Ifit2*^-/-^ CNS tissues. Myelin debris clearance is an important property of reparative M2 type myeloid cells, and we found that infiltrating myeloid cells and microglia express genes and metabolic profiles associated with proinflammatory M1 type myeloid cells in *Ifit2*^-/-^ mice. Therefore, these results demonstrate that Ifit2 may be a critical ISG that mediates IFN-I’s anti-inflammatory functions by supporting reparative M2 differentiation and functions, which promote myelin debris clearance and remyelination.

## Materials and Methods

### Mice

C57BL/6 (B6) mice were purchased from The Jackson Laboratory (Bar Harbor, ME). IFIT-WL KO and *Ifit2*^-/-^ mice were previously reported (19, 20). All mouse strains were housed, maintained, and bred under specific pathogen-free conditions at the Northwestern University Feinberg School of Medicine. All experimental procedures with mice were approved by the Institutional Animal Care and Use Committee (IACUC) of the Northwestern University and of the Cleveland Clinic Foundation.

### Induction of EAE

Mice were subcutaneously immunized on the back with 300μg of MOG_35-55_ peptide (BioSynthesis, Lewisville, TX) emulsified in equal volume of CFA containing 5 mg/ml Mycobacterium tuberculosis H37Ra (Difco, Detroit, MI). Mice were also injected with 200 ng pertussis toxin (Sigma, St. Louis, MO) intraperitoneally on day 0 and 2 post immunization. EAE development was monitored and graded on a scale of 0 to 5: 0, no disease; 1, limp tail; 2, hind limb weakness; 3, hind limb paralysis; 4, hind and fore limb paralysis; 5, moribund or death.

### Real-Time Quantitative PCR (qPCR)

Mice were euthanized at peak of disease and perfused with cold PBS. Total RNA was extracted from homogenized brains and spinal cords using TRIzol reagent according to the manufacturer’s instructions (Invitrogen). CD45^high^CD11b^high^ (infiltrating monocyte), CD45^int^ CD11b^high^ (microglia) and CD45^low^ (astrocyte and oligodendrocyte) cells were sorted from the CNS using a FACSMelody cell sorter (BD Bioscience) and RNA was extracted using a Quick-RNA Miniprep Kit (Zymo Research, Irvine, CA). cDNA was generated with MMLV reverse transcriptase (Promega, Madison, WI). Real-time PCR was performed using Radiant qPCR mastermix (Alkali Scientific, Fort Lauderdale, FL) or SYBR Green Master Mix (Applied Biosystems, Waltham, MA) in a QuantStudio 3 Real-Time PCR System (Applied Biosystems). All genes were normalized to *Gapdh*. The following SYBR primers and Taqman probes were listed in Table 1.

**Table 1.**
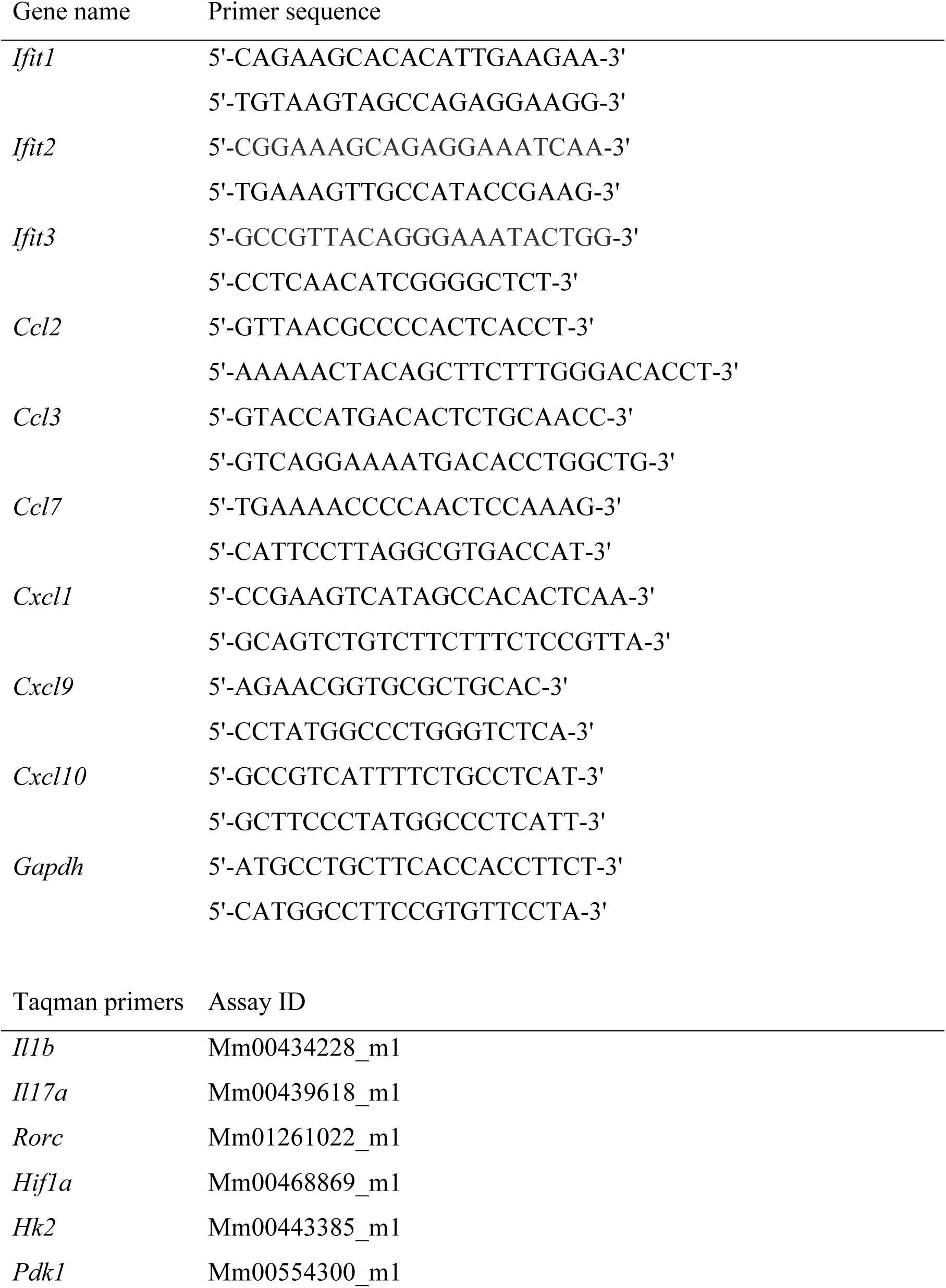
Primer sequences for RT-qPCR.

### Flow Cytometry

CNS mononuclear cells were isolated by 70%/30% Percoll density gradient. Surface staining of mononuclear cells was performed with anti-CD4 (RM4–5), anti-CD44 (IM7). anti-CD25 (PC61.5), anti-GITR (DTA-1), anti-ICOS (C398.4A), anti-CD11b (M1/70), anti-CD45 (I3/2.3), anti-MHC Class II (M5/114.15.2), anti-CD206 (C068C2) and anti-iNOS (CXNFT) antibodies. For intracellular cytokine detection, cells were stimulated for 4 h with PMA (10ng/mL, Millipore-Sigma) / ionomycin (1μM, Millipore-Sigma) for 4 h in the presence of 2μM monensin (Calbiochem) during the last 2 h of stimulation. Cells were then fixed with 4% paraformaldehyde, washed with PBS, permeabilized with perm buffer, and stained with anti-IL-17 (TC11-18H10), anti-IFNγ (XMG1.2), anti-TNFα (TN3-19), anti-GM-CSF (MP1-22E9) antibodies. The Abs were purchased from eBioscience (San Diego, CA), BD Biosciences (San Diego, CA) and BioLegend (San Diego, CA). Samples were acquired on a FACSCelesta (BD Biosciences) and analyzed with a FlowJo software (TreeStar, Ashland, OR).

### Immunohistochemistry of spinal cord sections

For immunohistochemistry, the WT and *Ifit2*^-/-^ mice were perfused intracardially with 4% paraformaldehyde (PFA) under anesthesia. The spinal cord was isolated and post-fixed with 4% PFA overnight and then placed in 30% sucrose at 4°C. After the samples had sunk, the spinal cord is dissected into cervical, thoracic and lumbar tissue blocks, embedded in OCT compound (Scigen-Tissue Plus) and frozen on dry ice. Cryostat sections (10μm) were cut on a Leica cryotome and affixed to Superfrost™ Plus Microscope Slides (FisherScientific). Sections were thawed, air-dried, rehydrated with PBS and further non-specific binding was blocked using 5% goat serum. The following antibodies were incubated in 0.1% Triton with 2% goat serum in PBS at 4°C overnight: rabbit anti-GFAP (1: 1000, DAKO, Z0334), rabbit anti-IBA-1 (1: 500, Abcam, AB178846), rabbit anti-TMEM119 (1:250, Abcam, ab209064), rat anti-MBP (1: 500, Millipore-Sigma, MAB386) and mouse anti-NF200 (1:500, Sigma, N5389) antibodies. Double staining was performed for MBP and GFAP, or IBA-1 and TMEM119). The sections were then washed in PBS and incubated with a secondary antibody conjugated to Alexa Fluor 488 or Alexa Fluor 555 (1:1000, Invitrogen) for 2 hrs. After washing, slides were mounted with prolong Gold antifade reagent with DAPI (Invitrogen). Images were taken with a Zeiss AX10 (Imager Z2) confocal microscope.

### Organotypic Slice culture

Organotypic cerebellar slice culture were established from 5 to 8-day-old pups from WT and *Ifit2*^-/-^ mice according to Tripathi et al. (21). Briefly, brains of P5 to P8 pups were removed and placed in ice-cold artificial cerebral spinal fluid (2 mM calcium chloride, 10 mM glucose, 3 mM potassium chloride, 4 mM magnesium sulphate, 26 mM sodium bicarbonate, 2 mM Ascorbic acid). Excess brain regions were removed and cerebellum was embedded in 2% low-melting agarose with artificial cerebrospinal fluid. Using Vibratome, 300 μm sagittal slices of the cerebellum were sectioned. Two to three slices were placed onto cell-culture inserts (Millicell 0.4m, Millipore) in media containing 50% MEM, 25% Horse Serum, 25% Hank’s Buffer, 1% GlutaMax, 10 mg/ml Glucose and 1x antibiotic-antimycotic. To study demyelination, slices, after five days in culture, were incubated with 0.5mg/ml lysolecithin for 20h. For remyelination, slices were washed with media (3x) and cultured an additional 6-7 days for remyelination. For immunofluorescence staining, slices were fixed with 4% PFA for 30 min, permeabilized with 2% Triton X-100 for 30 min, and blocked with 10% donkey serum and 0.1% Triton X-100 in PBS for 1 h at room temperature before being incubated in the primary antibodies against neurofilament (mouse anti-NF200, 1:750) and myelin (rat anti-MBP, 1:250) overnight at 4 °C. Primary antibodies were visualized by incubating sections with the appropriate Alexa fluorophore-conjugated secondary antibodies for 1h at room temperature. After mounting the slices, images were taken with a Zeiss AX10 (Imager Z2) confocal microscope.

### Statistical Analysis

Statistical significance was determined by the Mann-Whitney test using a Prism software (GraphPad, San Diego, CA). p<0.05 was considered statistically significant.

## Results

### Ifit expression is elevated during EAE

We first examined *Ifit* mRNA expression in the CNS tissues during EAE. C57BL/6 mice were immunized with MOG peptide in CFA followed by pertussis toxin administration to induce EAE. CNS tissues were harvested at disease onset (7 day), peak (day 14), and recovery (day 21) phases. Expression of all *Ifit* mRNAs (*Ifit1, Ifit2*, and *Ifit3*) in the CNS was determined by qPCR analysis. The expression pattern closely matched with the disease activity (Fig 1). It was previously reported that endogenous type I IFN production is localized in the target tissues during EAE and that the resulting enhanced IFNAR signaling activity is related to the disease severity (7). Since type I IFN is the primary stimulus inducing IFIT expression (22), increased *Ifit* mRNA expression in EAE is likely induced by IFN signaling elicited from the endogenously produced type I IFN. It is interesting to note that low but detectable basal level expression of *Ifit2* mRNA is observed in naïve CNS tissue (Fig 1).

**Fig 1.**
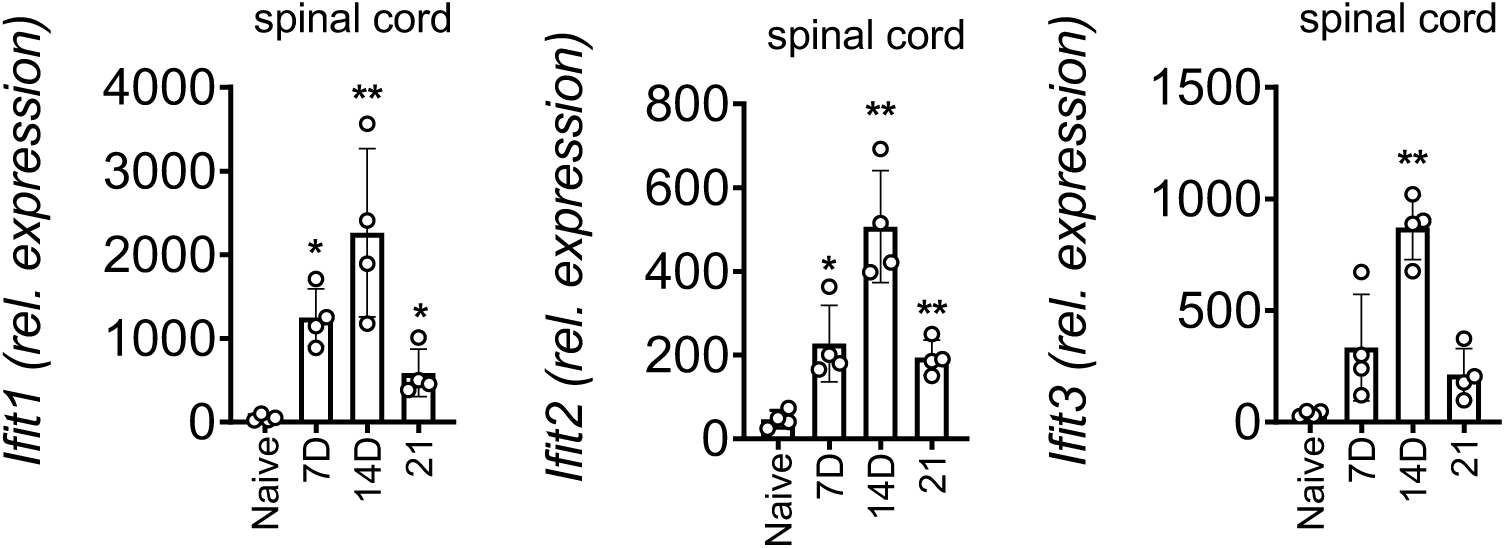
Expression of *Ifit* mRNA in the CNS during EAE. C57BL/6 mice were injected with CFA/MOG_35–55_ followed by pertussis toxin on the day of immunization and 48 h later to induce EAE. qPCR from whole spinal cord were harvested at disease onset (7 day), peak (day 14), and recovery (day 21) phases. Data were normalized to naive mice. n = 3-4 per group. *p < 0.05; **p < 0.01; as determined by Mann-Whitney nonparametric test.

### Ifit whole locus mutant mice are highly susceptible to EAE

We previously reported using Ifit whole locus (IFIT-WL) knockout mice that viral replication is drastically increased in the absence of the Ifit whole locus (19). To investigate if the lack of Ifit family proteins affects autoimmune inflammation, wild type and IFIT-WL KO mice were induced for EAE. The onset and initial progress appeared to be similar between the groups. However, IFIT-WL KO animals exhibited more severe disease based on the clinical score (Fig 2A). Consistent with the disease severity, the total numbers of CD4 T cells infiltrating the CNS tissue were significantly increased in KO mice, although the proportion was comparable (Fig 2B). Foxp3^+^ regulatory T (Treg) cells accumulated in the inflamed sites are important in resolving inflammation (23). The proportion of CNS infiltrating Treg cells was similar between the groups (Fig 2B); however, the absolute number of Treg cells was greater in KO mice. Surface expression of Treg cell associated markers, such as ICOS, GITR, and CD25, remained comparable (Fig 2C). These results suggest that the susceptibility seen in KO mice is not attributed to defects in Treg cell recruitment or activation. We then found that CNS accumulation of effector CD4 T cells expressing encephalitogenic cytokines, IL-17, TNFα, GM-CSF, and IFNγ, were markedly increased in both proportions and total numbers (Fig 2D). This was further supported by gene expression level shown in Supplementary Fig 1. None of the *Ifit* mRNA was found in IFIT-WL KO mice, while all three *Ifit* mRNAs were drastically increased in wild type CNS tissues (Supp Fig 1A). Encephalitogenic cytokine mRNA expression was consistent with flow cytometry data. *Il17a, Il1b, Tnfa*, and *Rorc* mRNA expression was markedly elevated in IFIT-WL KO mice compared to those in WT control mice (Supp Fig 1B). These results demonstrate that Ifit proteins play an important immune regulatory role in controlling encephalitogenic inflammatory responses.

**Fig 2.**
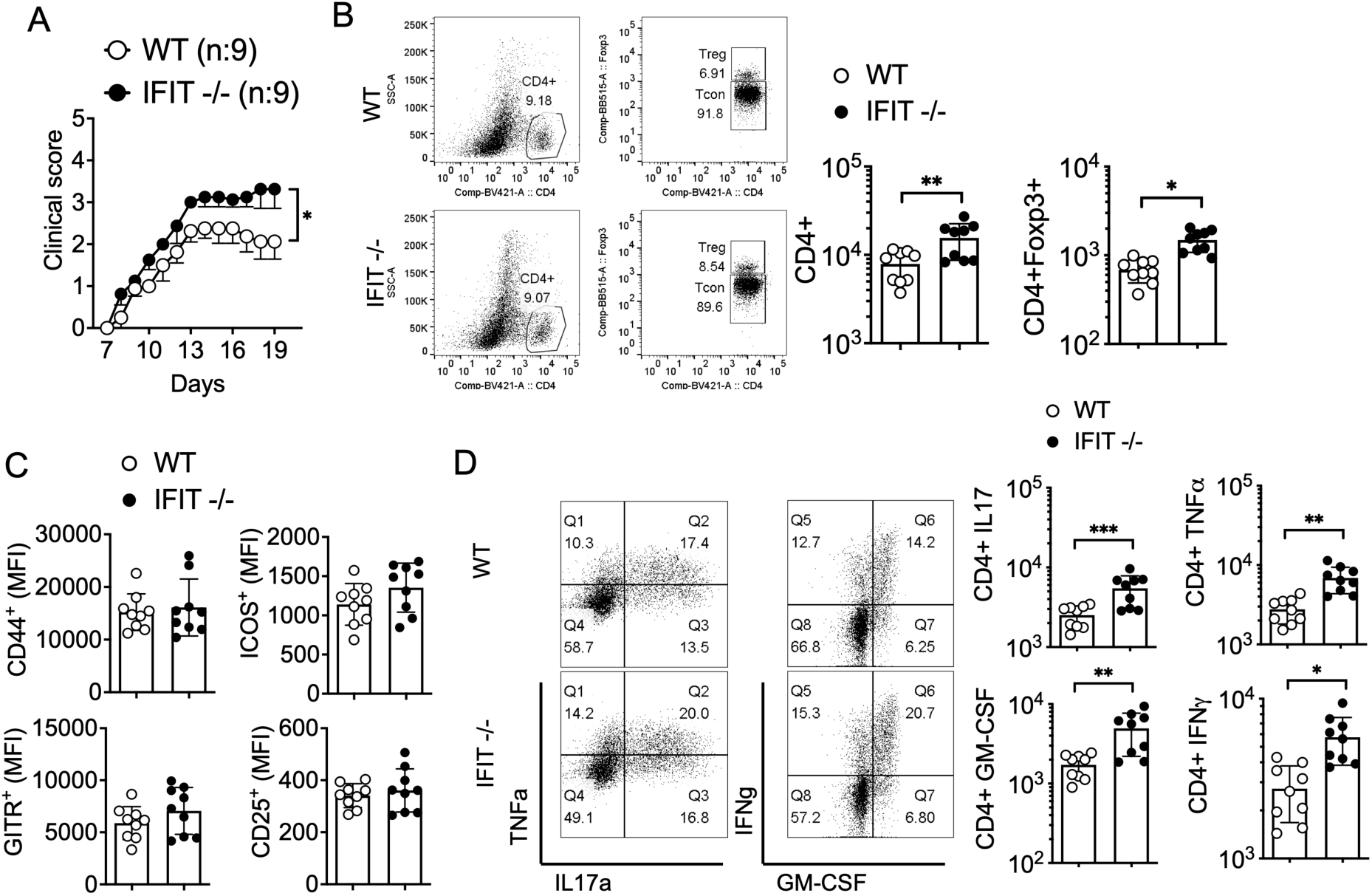
Increased susceptibility to EAE in IFIT-WL KO mice. WT (n = 9) and IFIT-WL KO (n = 9) mice induce for EAE. (**A**) Evaluated for clinical signs for 20 d. (**B**) Absolute number of CNS-infiltrating CD4^+^ T cells and CD4^+^Foxp3^+^ Treg cells in the CNS of WT and IFIT-WL KO mice at the peak of disease (day 17 post immunization). (**C**) The Mean Fluorescence Intensity (MFI) of CD44, ICOS, GITR and CD25 on CD4^+^Foxp3^+^ Treg cells in the CNS. (**D**) Total numbers of GM-CSF, IFNγ, IL-17, and TNFα expressing CD4^+^ T cells infiltrating the CNS at the peak of disease (day 17 post immunization). n = 9 per group. *p < 0.05; **p < 0.01; ***p < 0.001; as determined by Mann-Whitney nonparametric test.

### Ifit2 KO mice are highly susceptible to EAE

While the canonical functions of the Ifit proteins could be overlapping, there is evidence that each Ifit could still play a distinct role. For example, Ifit2 protects mice from lethal VSV induced neuropathogenesis, while Ifit1 protein is dispensable in anti-VSV immunity (24). Ifit2 also plays a unique role in protecting mice from coronavirus induced encephalitis by regulating microglial activation and leukocyte migration (16). Thus, we investigated if Ifit2 plays a role in EAE independent of other Ifit proteins. *Ifit2*^-/-^ mice were induced for EAE. Analogous to what we observed from IFIT-WL KO mice, we found that the disease onset and initial progress were comparable in WT and *Ifit2*^-/-^ mice. However, *Ifit2*^-/-^ mice failed to recover from the acute paralysis and continued to display severe disease as in IFIT-WL KO mice (Fig 3A). CD4 T cells infiltrating the CNS tissues were markedly increased in *Ifit2*^-/-^ mice, although the proportions of CD4 T cells seem comparable between the groups as seen in IFIT-WL KO mice (Fig 3B). Proportion of CNS infiltrating Treg cells were similar (data not shown); however, the absolute numbers were elevated in *Ifit2*^-/-^ mice (Fig 3B). Likewise, Treg cell marker expression such as ICOS, GITR, and CD25 remained unchanged (data not shown). We then examined proinflammatory cytokine expression profiles in CNS infiltrating CD4 T cells. As shown in Fig 3C, CD4 T cell expression of inflammatory cytokines was drastically increased in *Ifit2*^-/-^ mice. Consistent with escalated proinflammatory CD4 T cell profiles in the CNS, the expression of key transcription factors for Th1 and Th17 immunity, *Tbx21* and *Rorc*, was similarly increased in *Ifit2*^*-/-*^ mice (Fig 3D and data not shown). Therefore, Ifit2 alone may play an immune regulatory function to control the development of encephalitogenic immune responses.

**Fig 3.**
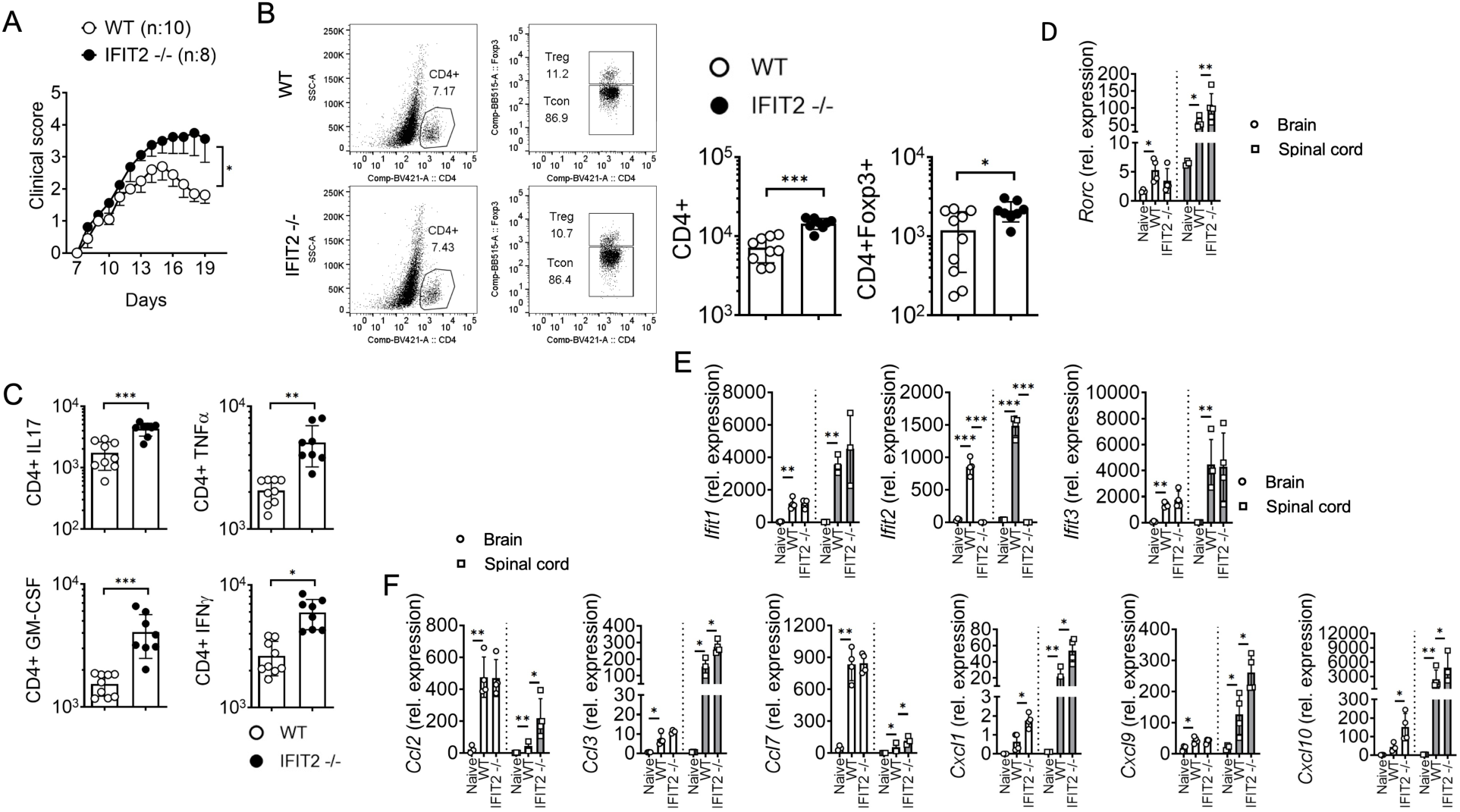
*Ifit2*^-/-^ mice were more susceptible to EAE than WT mice. WT (n = 10) and *Ifit2*^-/-^ (n = 8) mice induce for EAE. (**A**) Time course of the development of EAE. (**B**) Absolute number of CNS-infiltrating CD4^+^ T cells and CD4^+^Foxp3^+^ Treg cells in the CNS of WT and *Ifit2*^-/-^ mice at the peak of disease (day 17 post immunization). (**C**) Numbers of GM-CSF, IFNγ, IL-17, and TNFα expressing CD4^+^ T cells infiltrating the CNS at the peak of disease (day 17 post immunization). n = 8-9 per group. (**D-F**) qPCR analysis of the indicated mRNAs in the brain and spinal cords from naïve, WT, and *Ifit2 -/-* mice at the peak of disease (day 17 post immunization). n = 3 per group. *p < 0.05; **p < 0.01; ***p < 0.001; as determined by Mann-Whitney nonparametric test.

Ifit2 deficiency did not affect IFN-I expression in EAE (data not shown). In support, other Ifit, *Ifit1* and *Ifit3*, mRNA expression in the CNS tissues of *Ifit2*^*-/-*^ mice remained comparable to that of wild type mice, strongly suggesting that endogenous IFN-I production is not affected by the absence of Ifit2 (Fig 3E). This result also suggests that the susceptibility of *Ifit2*^-/-^ mice to EAE cannot be attributed to impaired IFN-I or other Ifit expression and that Ifit2 plays a unique role in modulating inflammatory responses.

Severe EAE and increased T cell infiltration in the CNS were further corroborated by enhanced inflammatory chemokine expression. CCL2 and CCL3 production in the CNS tissue critically regulates EAE pathogenesis through recruiting macrophages and dendritic cells (25, 26). Consistent with severe pathogenesis in *Ifit2*^*-/-*^ mice, we found that *Ccl2* and *Ccl3* mRNA expression particularly in the spinal cord was significantly elevated (Fig 3F). CCL7 is an alternative CCR2 ligand potentially involved in EAE pathogenesis (27, 28), and indeed the *Ccl7* expression was also found increased in the *Ifit2*^-/-^ spinal cord (Fig 3F). Overexpression of CXCL1 in astrocytes enhances EAE development by supporting inflammatory cell recruitment (29). Likewise, CXCR3 ligands, CXCL9 and CXCL10 have been implicated in EAE, where the expression is significantly increased in the CNS (30). Consistent with these reports, we found that Ifit2 deficiency markedly increased all those chemokines *Cxcl1, Cxcl9*, and *Cxcl10* (Fig 3F).

APC-derived IL-12 family cytokines are especially important in tuning the activation of pathogenic T cells such as Th17 type cells. We found that IL-23p19 subunit expression was dramatically increased in *Ifit2*^-/-^ mice, although IL-12p35 and IL-12p40 subunit expression remained unchanged (Supp Fig 2). IL-27, composed of IL-27p28 and EBI3 subunits and induced by IFN-I signaling, plays an important role in encephalitogenic inflammation (8, 31). While IL-27 could mediate protective roles in EAE pathogenesis, its expression also represents the extent of inflammatory activity. In support, we found elevated expression of both subunits of IL-27 in *Ifit2*^-/-^ mice (Supp Fig 2 and data not shown). Taken together, these results demonstrate that Ifit2 deficiency displays broad impacts on unleashing inflammatory mediator expression in the CNS tissues, supporting encephalitogenic immune responses.

### Altered phenotypes of myeloid cells in the CNS of Ifit2^-/-^ mice

An imbalance between demyelination and remyelination could trigger disease process in multiple sclerosis, and clearing myelin debris derived from demyelination is an important process promoting remyelination (32). Myeloid cells including monocyte-derived macrophages or microglia are the central cell types mediating myelin debris clearance (32). Macrophages and microglia undergo differentiation into two distinct subsets, proinflammatory M1 and regulatory M2 phenotype cells, and M2 phenotype cells are specifically known to induce reparative processes (33). In case of demyelinating inflammation, it was reported that M2 phenotype microglia remove myelin debris from the lesion, further promoting remyelination process (34, 35). We thus compared infiltrating myeloid cells and CNS resident microglial cells. Infiltrating myeloid cells defined as CD45^high^CD11b^high^ cells were highly abundant in *Ifit2*^-/-^ mice (Fig 4A). Their MHCII expression was significantly higher in *Ifit2*^-/-^ mice. Importantly, expression of iNOS, a marker for M1 type pro-inflammatory subset, was also higher in *Ifit2*^-/-^ cells. On the other hand, expression of the mannose receptor, CD206, a marker for M2 type anti-inflammatory subset, was substantially lower in *Ifit2*^-/-^ cells (Fig 4A). Microglia, tissue resident macrophages in the CNS, have also been implicated in neuroinflammation (36), and are important sources of cytokines and chemokines. They can similarly polarize into M1 or M2 phenotype cells. In EAE, microglia undergo expansion and M1 type differentiation (37, 38). In support, we observed greater expansion of microglia population (CD45^int^CD11b^high^) in *Ifit2*^-/-^ mice (Fig 4B). Likewise, microglia expression of MHCII molecule was drastically elevated in *Ifit2*^-/-^ mice. Unlike infiltrating myeloid cells, upregulation of M1-like marker, iNOS, was not observed. However, downregulation of CD206 was pronounced (Fig 4B). Therefore, Ifit2 deficiency appears to have significant impact on myeloid cells and microglia activation and polarization, possibly contributing the pathogenesis.

**Fig 4.**
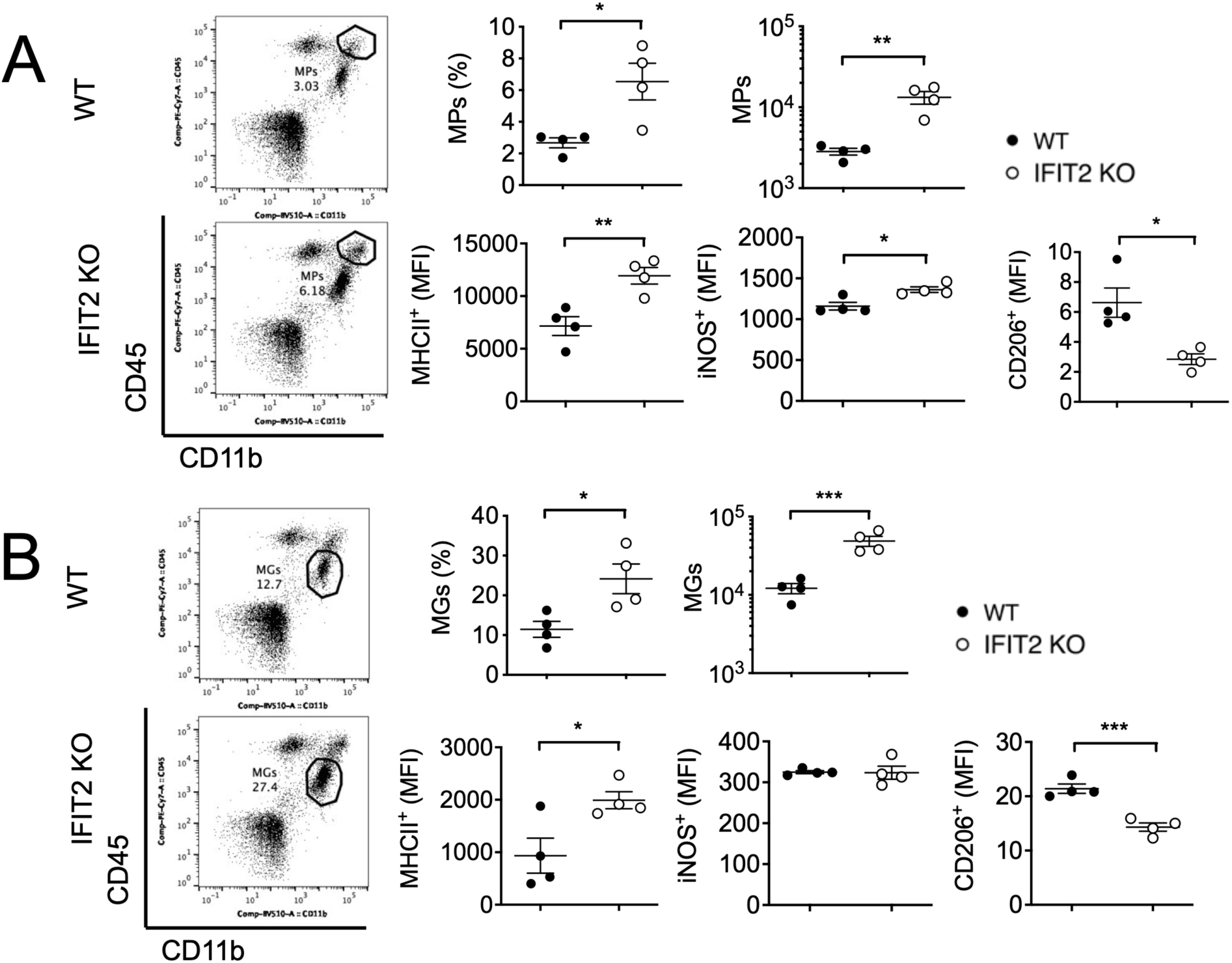
Phenotypes of myeloid subsets in the CNS of *Ifit2*^*-/-*^ mice. Representative flow cytometry plots of infiltrating myeloid cells (**A**) and of microglia (**B**) from CNS of WT and *Ifit2 -/-* mice. Absolute numbers of myeloid cells (CD45^hi^CD11b^hi^) and microglia (CD45^int^ CD11b^hi^). The Median Fluorescence Intensity (MFI) of MHCII, iNOS and CD206 expression measured by flow cytometry. n = 4 per group. *p < 0.05; **p < 0.01; ***p < 0.001; as determined by Mann-Whitney nonparametric test.

### Ifit2 deficiency exacerbates demyelination and axonal loss in EAE

We next investigated if Ifit2 deficiency alters demyelination/remyelination during autoimmune inflammation of EAE. Immunohistochemistry of coronal spinal cord sections from WT and *Ifit2*^- /-^ mice with EAE revealed significant changes in myelination and axonal density in the cervical, thoracic, and lumber spinal cords. As seen by lower MBP staining in spinal cord sections, EAE generated a notable myelin damage in *Ifit2*^-/-^ relative to WT at the peak of disease (Supp Fig 3A). The ventral white matter of the thoracic area of the *Ifit2*^-/-^ mice spinal cord showed widespread and notable myelin degradation. In the WT and *Ifit2*^-/-^ spinal cords, we performed double immunostaining with antibodies to MBP and NF200 to determine the condition of demyelinated axons and found a rather intact myelin ring (green) around the axons (red). Interestingly, we found several MBP^+^ myelin sheaths in *Ifit2*^-/-^ mice that were devoid of NF200 labeled axons (Supp Fig 3A, arrows), implying that axons have been destroyed or damaged and are consequently negative for NF200 antibody.

### Ifit2 deficiency exacerbates microgliosis, macrophage infiltration and astrocyte activation in EAE

Microglia, macrophages, and astrocytic activation are all required for immune-mediated demyelination. We looked at whether the spinal cords of MOG 35-55 immunized WT and *Ifit2*^-/-^ mice revealed any differences in peripheral macrophage infiltration and/or microglial activation to see if greater microgliosis/ macrophage activity was associated with severe EAE-related pathology in *Ifit2*^-/-^ animals. In the cervical, thoracic, and lumbar areas of the WT and *Ifit2*^-/-^ mice spinal cords, immunohistochemistry for ionized calcium binding adaptor molecule-1 (IBA-1) was used to mark macrophage/microglia. The density of IBA-1^+^ cells in the ventral white matter of all regions of the spinal cord was significantly greater in *Ifit2*^-/-^ mice than in WT mice, indicating more immune cell infiltration in *Ifit2*^-/-^ mice (Fig 5A). MBP^+^ myelin debris are engulfed by IBA^+^ macrophages, and immune cell infiltration and phagocytosis are more evident in the thoracic area (Fig 5B), according to our findings. Of note, infection with the neurotrophic coronavirus MHV-RSA59 resulted in decreased microglial activation and infiltration of peripheral immune cells (NK, CD4 T cells) into the brain in *Ifit2*^-/-^ mice (39), suggesting a unique mode of pathogenesis of EAE in the absence of Ifit2. Microglia and peripheral macrophages have different origins, and microglia are activated prior to peripheral immune infiltration and actively assist in their recruitment, which may also be true in EAE. We stained the spinal cord sections for transmembrane protein 119 (TMEM119), which has emerged as a microglia-specific marker, to assess the alteration in microglial numbers. At the peak of disease, TMEM119 staining revealed that *Ifit2*^-/-^ mice had a higher number of microglia than WT mice, supporting the flow cytometry data shown in Fig 4B. *Ifit2*^-/-^ mice’s TMEM119^+^ microglia have an active phenotype with amoeboid morphology and retracted microglial processes (Fig 5C). Several reports have showed that astrocyte regulate CNS inflammation by affecting infiltration of peripheral immune cells and activation of resident microglia (40, 41). We next determined astrocyte activation by immunohistochemistry using antibody against GFAP. We observed that *Ifit2*^-/-^ mice show higher GFAP^+^ astrocytes compared to WT at the peak of EAE. The degree of astrocyte reactivity is positively correlated with GFAP expression (42, 43), hence indicates more astrocyte activation in *Ifit2*^-/-^ mice.

**Fig 5.**
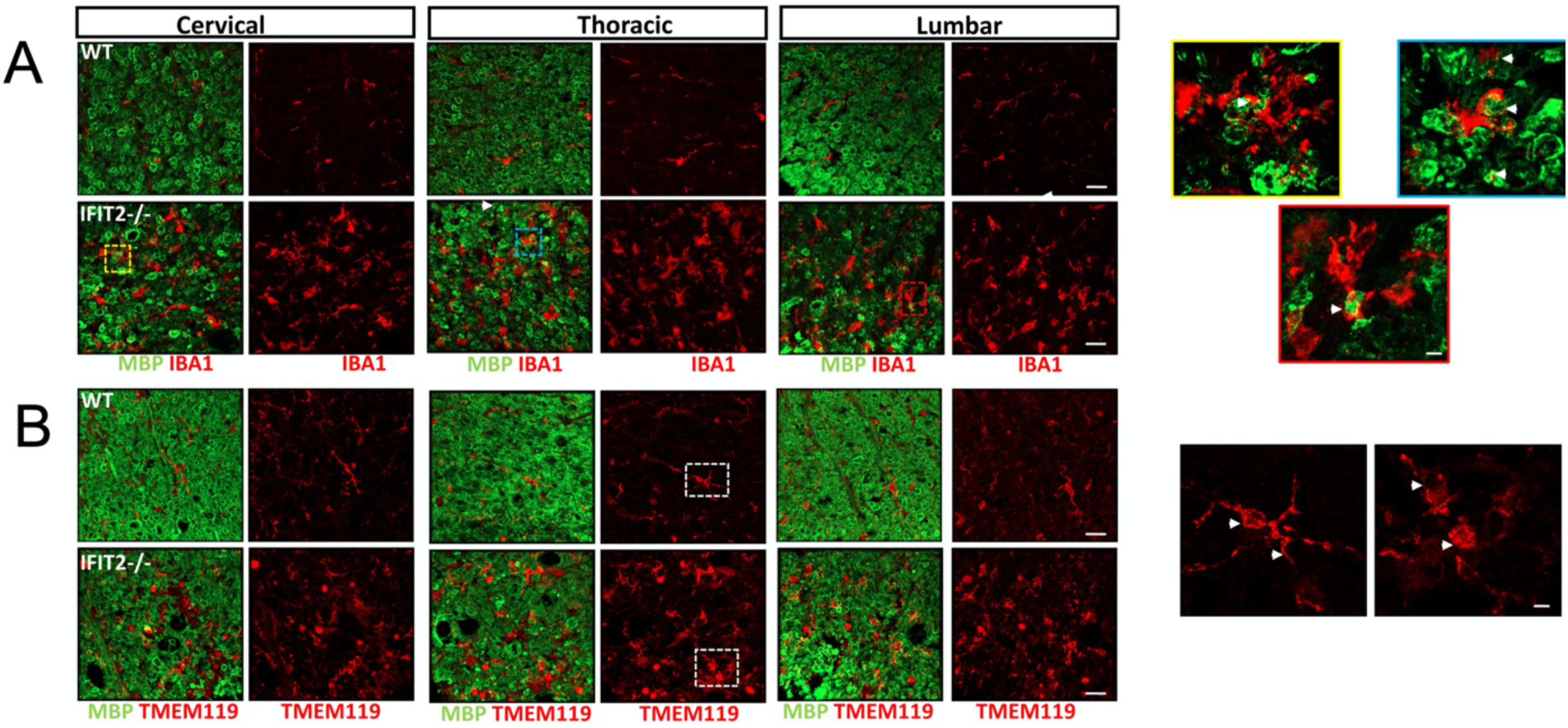
Ifit2 deficiency augments number of macrophage/microglia following EAE induction at the peak of disease. Immunohistochemical staining of macrophage/microglia cell types in ventral white matter of cervical, thoracic, and lumbar spinal cord of WT and *Ifit2*^-/-^ mice. (**A**) IBA-1 macrophage/microglia marker of inflammation (Scale bar:20μm). Inset images showing engulfment of myelin debris by IBA1^+^ macrophage/microglia in cervical, thoracic, and lumbar regions (arrow, Scale bar: 5 μm) (**B**) TMEM119 staining, a marker of CNS local microglia in the ventral white matter of cervical, thoracic, and lumbar spinal cord of WT and *Ifit2*^-/-^ (Scale bar: 20μm). Insets display examples of microglial cells in WT (ramified, long branched processes) and *Ifit2*^-/-^ (amoeboid, round) mice (Scale bar: 5 μm).

### Greater demyelination, lack of myelin debris clearance, and loss of remyelination efficiency in Ifit2^-/-^ CNS tissues

M2 type microglia and macrophages support CNS remyelination (34). Since both infiltrating monocytes and microglia expressing M2 phenotype (CD206) were diminished in *Ifit2*^-/-^ mice, we sought to test if such defects in M2 type subset differentiation are associated with impaired remyelination and with severe pathogenesis seen in *Ifit2*^-/-^ mice. We used the model for organotypic slice cultures and lysolecithin (LPC, 0.5mg/ml) induced demyelination and remyelination in P10 control and *Ifit2*^-/-^ brains using our recent protocols (21, 44). Slices stained for myelin protein MBP and axonal marker NF200 show demyelination by LPC (Fig 6C) compared to control (Fig 6A). The extent of demyelination was, however, strikingly pronounced in slices from *Ifit2*^-/-^ brains (Fig 6G). Of particular interest was the lack of myelin debris clearance in the *Ifit2*^-/-^ brains (Fig 6G, shown by the yellow outline). We then examined the extent of remyelination as the lack of myelin clearance by the microglia and/or macrophages could have negative impact on remyelination (32). Indeed, spontaneous remyelination was observed in control slices (Fig 6E), while remyelination was severely impaired in *Ifit2*^-/-^ brains (Fig 6I). Of note, Ifit2 deficiency did not induce axonal loss (Fig 6D, 6F, 6H, and 6J), suggesting defects in remyelination in *Ifit2*^-/-^ brains. As the overall number of OL lineage cells were comparable in control and *Ifit2*^-/-^ mouse brains (data not shown), defective remyelination appears to be attributed to impaired myelin debris clearance.

**Fig 6.**
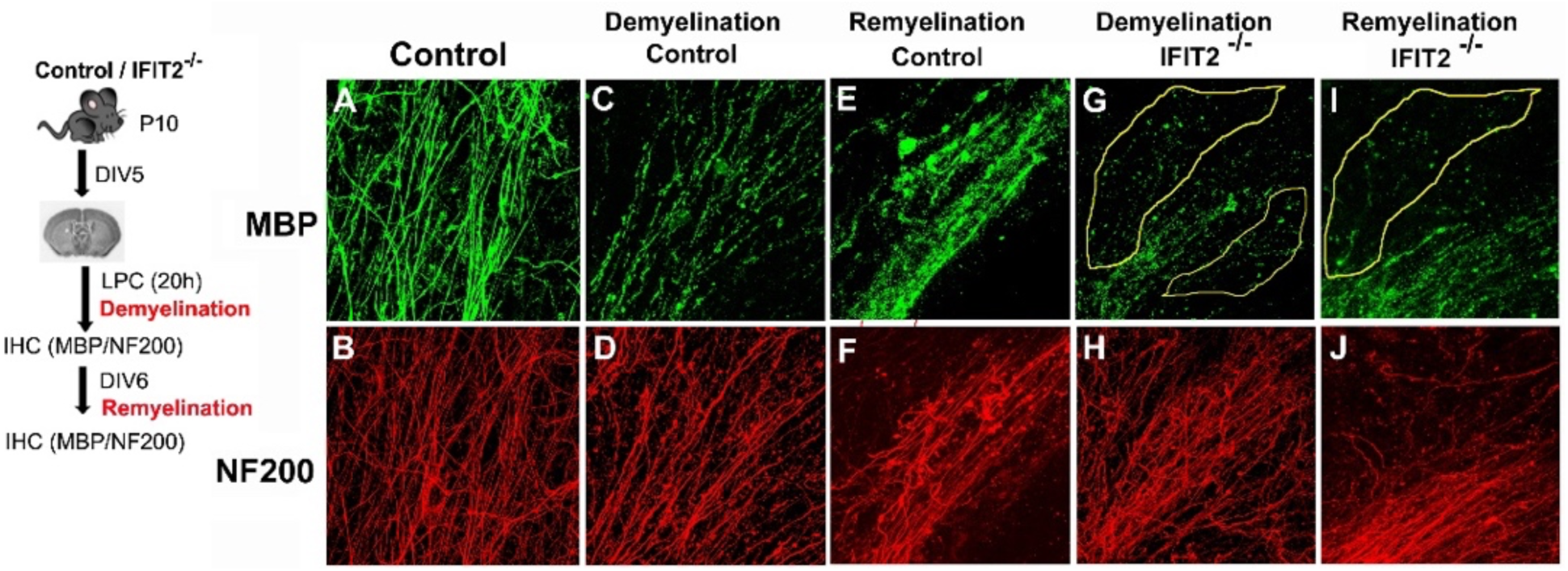
Ifit2 deletion is associated with increased demyelination, failed myelin debris clearance, and impaired remyelination. Mouse brain organotypic sections from control mice stained for myelin basic protein (MBP) and axonal marker NF200. Organotypic slices from control and *Ifit2*^-/-^ P10 mice were treated with lysolecithin (LPC) (0.5mg/ml) for 18 hrs to induce demyelination. Compared to control slices (**C**,**D**), *Ifit2*^-/-^ tissue section showed severe demyelination (**G**,**H**). Punctate myelin debris (**G**) were found in demyelinated areas in *Ifit2*^-/-^ brain slices (yellow borders; **G**). There was comparable number of axons between the two conditions (**D**,**H**). Following demyelination, the slices were allowed to recover and remyelinate for 7 days. Similar staining was carried out. Control slices (**E**,**F**) had significant recovery and remyelination. *Ifit2*^-/-^ brain slices had drastically lower extent of remyelination (**I-J**) despite almost similar levels of axons between conditions (**F**,**J**). There was myelin debris found during remyelination (yellow border; **I**).

### Ifit2 deficiency results in metabolic dysregulation of myeloid cells

Altered M1/M2 phenotypes in infiltrating myeloid cells and in microglia (Fig 4), exacerbated demyelination (Fig 5), and defects in myelin debris clearance and remyelination observed in *Ifit2*^-/-^ brain tissues prompted us to test if Ifit2 deficiency dysregulates metabolic programming of those myeloid cells. Notably, inflammatory M1 phenotype cells predominantly utilize glycolytic pathways as an energy source, whereas reparative M2 phenotype cells preferentially utilize oxidative phosphorylation (45, 46). We first measured proinflammatory and metabolic gene expression in WT and *Ifit2*^-/-^ bone marrow derived macrophages (BMDM) following stimulation with various stimuli, including LPS, IFNγ, and IL-4 to promote M1 and M2 differentiation, respectively. We found that *Ifit2*^-/-^ BMDM expressed greater proinflammatory cytokines, *Tnfa* and *Il1b*, following LPS stimulation. In addition, glycolytic gene, *Hif1a* and *Hk2* expression was also elevated in *Ifit2*^-/-^ BMDM cells (Fig 7A). Interestingly, IFNγ stimulation dramatically enhanced *Il1b* mRNA expression in *Ifit2*^-/-^ BMDM cells, although other tested genes remained unchanged. IL-4 stimulation, which promotes M2 macrophage differentiation (47), also strikingly enhanced *Il1b* and *Hif1a* expression in *Ifit2*^-/-^ BMDM cells (Fig 7A). Therefore, Ifit2 deficiency is associated with dysregulated inflammatory cytokine and metabolic gene expression in myeloid cells.

**Fig 7.**
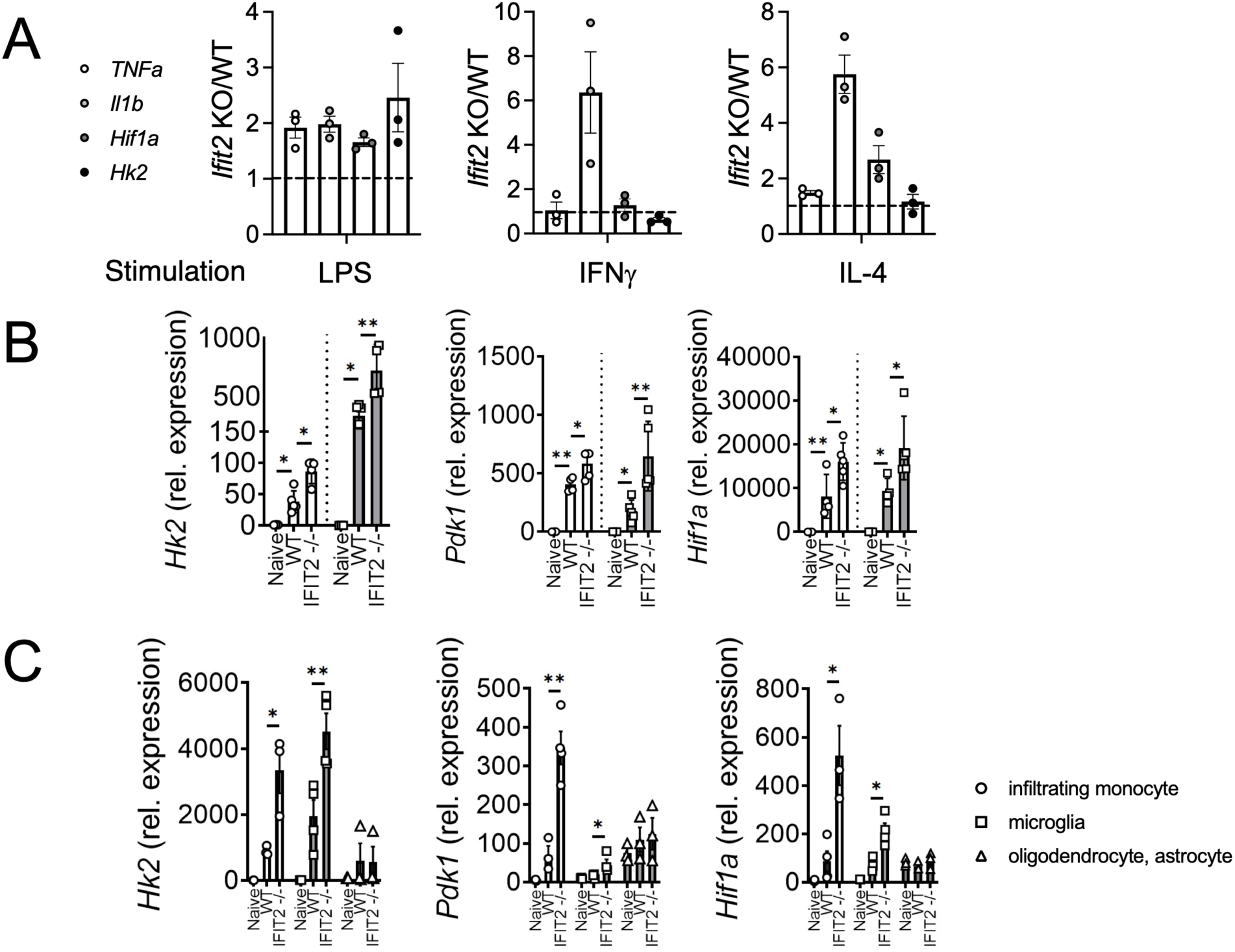
Cytokines and glycolytic gene expression in *Ifit2*^*-/-*^ mice. (**A**) mRNA expression of *Tnfa, Il1b, Hif1a, and Hk2* in bone marrow-derived macrophages from WT and *Ifit2*^*-/-*^ mice were incubated with lipopolysaccharide (LPS; 1 μg/mL) or IFNγ (20ng/ml) or IL-4 (5000U/mL) for 48 h. n = 3 per group. (**B**) CNS tissues harvested at the peak of disease were measured for glycolytic gene (*Hk2, Pdk1*, and *Hif1a*) expression by qPCR. (**C**) Sorted CD45^high^ CD11b^high^ (infiltrating monocyte), CD45^int^ CD11b^high^ (microglia) and CD45^low^ (astrocyte and oligodendrocyte) cells from CNS of EAE induced mice (day 17 post immunization) and the indicated gene expression were measured by qRT-PCR. n = 3-4 per group. *p < 0.05; **p < 0.01; as determined by Mann-Whitney nonparametric test.

To further investigate metabolic program in vivo, we isolated infiltrating myeloid cells (CD45^high^ CD11b^high^), microglia (CD45^int^ CD11b^high^), and CD45^low^ cells (including oligodendrocytes and astrocytes) from wild type and *Ifit2*^-/-^ mice induced for EAE. Genes involved in glycolytic pathway were measured. As shown in Fig 7B, expression of *Hk2, Pdk1*, and *Hif1a* was strikingly enhanced in *Ifit2*^-/-^ mice compared to WT mice. Of note, the enhanced gene expression was most pronounced in infiltrating myeloid cells and microglia. Therefore, these results strongly suggest that Ifit2 may control metabolic programming of myeloid cells via M1/M2 type differentiation, which may then have direct impact on disease progression, particularly remyelination process through myelin debris clearance.

## Discussion

In the present study, we report an unexpected role of Ifit2, one of the type I IFN-induced gene product known for the anti-viral functions, in modulating autoimmune inflammation. Mice deficient in the *Ifit2* gene were highly susceptible to myelin antigen-reactive autoimmunity, EAE. The susceptibility was closely associated with heightened encephalitogenic T cell responses that were accompanied by elevated proinflammatory cytokine and chemokine production. Importantly, we noticed that the lack of Ifit2 resulted in severe demyelination and impaired remyelination process, which appear to contribute to the exacerbated disease. Myelin debris clearance predominantly mediated by reparative macrophage and microglia subset is known to play a key role in supporting remyelination. One notable feature of proinflammatory and tissue repair subsets of myeloid cells is their metabolic program, i.e., glycolytic pathways in proinflammatory M1 type cells vs. oxidative phosphorylation in reparative M2 type cells. Indeed, we found that, in *Ifit2*^-/-^ infiltrating myeloid and CNS resident microglia, expression of genes involved in glycolytic pathways is markedly increased. Therefore, our results demonstrate that Ifit2 plays a novel regulatory function limiting inflammatory responses, in part by balancing metabolic profiles of myeloid lineage cells.

To our knowledge, the present study reports for the first time that Ifit2 expresses a novel immune regulatory function in autoimmunity. In EAE, all three Ifit family proteins (Ifit1, Ifit2, and Ifit3) are strongly induced in the spinal cord, and the kinetics of the expression closely mirror the disease activity. Given that endogenous IFN-I production in EAE is predominantly found in the target CNS tissues (7), Ifits induced by endogenous IFN-I are also likely localized in the CNS. The entire Ifit locus knockout animals display exacerbated disease, and Ifit2 deficiency alone results in similar susceptibility to EAE, suggesting that Ifit2’s ability to regulate autoimmune inflammation may be unique to Ifit2 protein. In support, IFN-I, Ifit1, and Ifit3 expression in *Ifit2*^-/-^ mice remain unchanged, strongly suggesting that high susceptibility of *Ifit2*^-/-^ mice to EAE is not due to defective IFN-I signaling or other Ifit protein production.

Mice deficient in Ifnar are highly susceptible to EAE induction, suggesting that IFN-I stimulation and subsequent induction of Ifit2 are crucial in limiting inflammatory responses (7). Likewise, the lack of Trif, an adaptor molecule crucial to induce IFN-I production, results in exacerbated EAE (8). The precise upstream signal that activates TRIF to induce IFN-I production within the CNS remains to be determined. Protective functions of TRIF suggest that ligands activating TLR3 or TLR4 may be involved in inducing IFN-I production in EAE. In support, TLR3 stimulation was previously shown to suppress EAE (48). Moreover, neonatal exposure to LPS can suppress EAE by promoting tolerogenic dendritic cells (49). However, pretreatment of LPS only prior to EAE induction is known to delay onset of the disease, while the severity remains unchanged (50). Paradoxically, mice deficient in Tlr4 surprisingly develop more severe EAE, suggesting the pathogenic role (51). Alarmin S100A9 is an alternative ligand for TLR4 (52), and S100A9-deficient mice were previously shown to develop exacerbated EAE, suggesting protective roles (53). It will be important to precisely determine endogenous signals capable of activating the TRIF-IFN-I-IFIT signaling pathways in EAE pathogenesis.

Cell type specific Ifnar-deficient mouse models were previously used to determine the target cells responsible for the severe EAE seen in *Ifnar*^-/-^ mice (7). T cell- or brain-expression of Ifnar is found dispensable, while targeting the Ifnar in myeloid lineage cells phenocopies global IFNAR deficiency, suggesting that resident and inflammatory monocytes as well as CNS resident microglia are likely target cells of IFN-I (7). Based on these findings, we speculate that myeloid lineage cells are likely the source of Ifit2-expressing cells. Our recent report similarly identified a unique function of Ifit2 that protects the host from lethal neurotropic MHV infection by modulating microglia activation and recruitment (16). The exact mechanisms by which IFN-I is endogenously produced (i.e., via TRIF-dependent mechanism) and acts on the target cells (i.e., via IFNAR-IFIT2-dependent mechanism) to mediate protective roles will require further investigation.

It is interesting to note that Ifit2 deficiency is closely associated with severe demyelination and defective remyelination. Following demyelination, macrophages and microglia play a crucial role in clearing myelin debris, which is known to be instrumental to support for the subsequent remyelination to occur (32). From the LPC-induced demyelination/remyelination organotypic slice culture and immunofluorescence microscopic examination experiments, we observed that Ifit2 deficiency is associated with defective myelin debris clearance and remyelination processes. Importantly, myelin debris clearance is one key property of M2-type macrophage/microglia (32). Therefore, it is possible that Ifit2 deficiency may impair transition from proinflammatory M1 phenotype cells to reparative M2 phenotype cell differentiation, which occurs during EAE (34, 54). M1 and M2 type macrophages (and microglia) utilize different metabolic programs, glycolysis and oxidative phosphorylation, respectively. Indeed, we noticed elevated glycolytic gene expression in *Ifit2*^-/-^ BMDM. In addition, *Ifit2*^-/-^ BMDM expression of proinflammatory cytokines, especially IL-1b, was markedly increased compared to wild type cells, even after IL-4 stimulation, which is known to promote M2 differentiation (47). Consistent with this, IFN-I promotes fatty acid oxidation and oxidative phosphorylation (55). Therefore, we would argue that Ifit2 may be an important switch that balances metabolic programming of macrophages (and microglia) to trigger remyelinating tissue repair processes by supporting M2 differentiation. While IFN-I is a widely used therapeutic option in MS patients, there is a cohort of patient refractory to the treatment (56). Better defining the precise cellular and molecular mechanisms will be a subject of great importance to identify novel therapeutic approaches for both IFN-I-responsive and -refractory MS patients.

## Notes

Supported by: NIH grants AI125247 and CA068782.

Conflict of interest: The authors have no conflict of interest.

### Competing Interest Statement

The authors have declared no competing interest.

## Bibliography

1. Benveniste, E. N., and H. Qin. 2007. Type I interferons as anti-inflammatory mediators. Sci STKE 2007: pe70.

2. Chen, K., J. Liu, and X. Cao. 2017. Regulation of type I interferon signaling in immunity and inflammation: A comprehensive review. J Autoimmun 83: 1–11.

3. Sormani, M. P., and N. De Stefano. 2013. Defining and scoring response to IFN-beta in multiple sclerosis. Nat Rev Neurol 9: 504–512.

4. Filippini, G., L. Munari, B. Incorvaia, G. C. Ebers, C. Polman, R. D’Amico, and G. P. Rice. 2003. Interferons in relapsing remitting multiple sclerosis: a systematic review. Lancet 361: 545–552.

5. Inoue, M., K. L. Williams, T. Oliver, P. Vandenabeele, J. V. Rajan, E. A. Miao, and M. L. Shinohara. 2012. Interferon-beta therapy against EAE is effective only when development of the disease depends on the NLRP3 inflammasome. Sci Signal 5: ra38. PMC3509177

6. Shinohara, M. L., J. H. Kim, V. A. Garcia, and H. Cantor. 2008. Engagement of the type I interferon receptor on dendritic cells inhibits T helper 17 cell development: role of intracellular osteopontin. Immunity 29: 68–78. PMC2625293

7. Prinz, M., H. Schmidt, A. Mildner, K. P. Knobeloch, U. K. Hanisch, J. Raasch, D. Merkler, C. Detje, I. Gutcher, J. Mages, R. Lang, R. Martin, R. Gold, B. Becher, W. Bruck, and U. Kalinke. 2008. Distinct and nonredundant in vivo functions of IFNAR on myeloid cells limit autoimmunity in the central nervous system. Immunity 28: 675–686.

8. Guo, B., E. Y. Chang, and G. Cheng. 2008. The type I IFN induction pathway constrains Th17-mediated autoimmune inflammation in mice. J Clin Invest 118: 1680–1690. PMC2276397

9. Lee, A. J., and A. A. Ashkar. 2018. The Dual Nature of Type I and Type II Interferons. Front Immunol 9: 2061. PMC6141705

10. McNab, F., K. Mayer-Barber, A. Sher, A. Wack, and A. O’Garra. 2015. Type I interferons in infectious disease. Nat Rev Immunol 15: 87–103. PMC7162685

11. Pidugu, V. K., H. B. Pidugu, M. M. Wu, C. J. Liu, and T. C. Lee. 2019. Emerging Functions of Human IFIT Proteins in Cancer. Front Mol Biosci 6: 148. PMC6930875

12. Diamond, M. S., and M. Farzan. 2013. The broad-spectrum antiviral functions of IFIT and IFITM proteins. Nat Rev Immunol 13: 46–57. PMC3773942

13. Johnson, B., L. A. VanBlargan, W. Xu, J. P. White, C. Shan, P. Y. Shi, R. Zhang, J. Adhikari, M. L. Gross, D. W. Leung, M. S. Diamond, and G. K. Amarasinghe. 2018. Human IFIT3 Modulates IFIT1 RNA Binding Specificity and Protein Stability. Immunity 48: 487–499 e485. PMC6251713

14. Pichlmair, A., C. Lassnig, C. A. Eberle, M. W. Gorna, C. L. Baumann, T. R. Burkard, T. Burckstummer, A. Stefanovic, S. Krieger, K. L. Bennett, T. Rulicke, F. Weber, J. Colinge, M. Muller, and G. Superti-Furga. 2011. IFIT1 is an antiviral protein that recognizes 5’-triphosphate RNA. Nat Immunol 12: 624–630.

15. Yang, Z., H. Liang, Q. Zhou, Y. Li, H. Chen, W. Ye, D. Chen, J. Fleming, H. Shu, and Y. Liu. 2012. Crystal structure of ISG54 reveals a novel RNA binding structure and potential functional mechanisms. Cell Res 22: 1328–1338. PMC3434343

16. Das Sarma, J., A. Burrows, P. Rayman, M. H. Hwang, S. Kundu, N. Sharma, C. Bergmann, and G. C. Sen. 2020. Ifit2 deficiency restricts microglial activation and leukocyte migration following murine coronavirus (m-CoV) CNS infection. PLoS Pathog 16: e1009034. PMC7738193

17. John, S. P., J. Sun, R. J. Carlson, B. Cao, C. J. Bradfield, J. Song, M. Smelkinson, and I. D. C. Fraser. 2018. IFIT1 Exerts Opposing Regulatory Effects on the Inflammatory and Interferon Gene Programs in LPS-Activated Human Macrophages. Cell Rep 25: 95–106 e106. PMC6492923

18. Imaizumi, T., N. Sassa, S. Kawaguchi, T. Matsumiya, H. Yoshida, K. Seya, T. Shiratori, K. Hirono, and H. Tanaka. 2018. Interferon-stimulated gene 60 (ISG60) constitutes a negative feedback loop in the downstream of TLR3 signaling in hCMEC/D3 cells. J Neuroimmunol 324: 16–21.

19. Kimura, T., C. T. Flynn, M. Alirezaei, G. C. Sen, and J. L. Whitton. 2019. Biphasic and cardiomyocyte-specific IFIT activity protects cardiomyocytes from enteroviral infection. PLoS Pathog 15: e1007674. PMC6453442

20. Butchi, N. B., D. R. Hinton, S. A. Stohlman, P. Kapil, V. Fensterl, G. C. Sen, and C. C. Bergmann. 2014. Ifit2 deficiency results in uncontrolled neurotropic coronavirus replication and enhanced encephalitis via impaired alpha/beta interferon induction in macrophages. J Virol 88: 1051–1064. PMC3911674

21. Tripathi, A., C. Volsko, J. P. Garcia, E. Agirre, K. C. Allan, P. J. Tesar, B. D. Trapp, G. Castelo-Branco, F. J. Sim, and R. Dutta. 2019. Oligodendrocyte Intrinsic miR-27a Controls Myelination and Remyelination. Cell Rep 29: 904–919 e909. PMC6874400

22. Zhou, X., J. J. Michal, L. Zhang, B. Ding, J. K. Lunney, B. Liu, and Z. Jiang. 2013. Interferon induced IFIT family genes in host antiviral defense. Int J Biol Sci 9: 200–208. PMC3584916

23. Gilroy, D., and R. De Maeyer. 2015. New insights into the resolution of inflammation. Semin Immunol 27: 161–168.

24. Fensterl, V., J. L. Wetzel, S. Ramachandran, T. Ogino, S. A. Stohlman, C. C. Bergmann, M. S. Diamond, H. W. Virgin, and G. C. Sen. 2012. Interferon-induced Ifit2/ISG54 protects mice from lethal VSV neuropathogenesis. PLoS Pathog 8: e1002712. PMC3355090

25. Dogan, R. N., A. Elhofy, and W. J. Karpus. 2008. Production of CCL2 by central nervous system cells regulates development of murine experimental autoimmune encephalomyelitis through the recruitment of TNF-and iNOS-expressing macrophages and myeloid dendritic cells. J Immunol 180: 7376–7384.

26. Manczak, M., S. Jiang, B. Orzechowska, and G. Adamus. 2002. Crucial role of CCL3/MIP-1alpha in the recurrence of autoimmune anterior uveitis induced with myelin basic protein in Lewis rats. J Autoimmun 18: 259–270.

27. Mildner, A., M. Mack, H. Schmidt, W. Bruck, M. Djukic, M. D. Zabel, A. Hille, J. Priller, and M. Prinz. 2009. CCR2+Ly-6Chi monocytes are crucial for the effector phase of autoimmunity in the central nervous system. Brain 132: 2487–2500.

28. Zhang, Y., J. J. Han, X. Y. Liang, L. Zhao, F. Zhang, J. Rasouli, Z. Z. Wang, G. X. Zhang, and X. Li. 2018. miR-23b Suppresses Leukocyte Migration and Pathogenesis of Experimental Autoimmune Encephalomyelitis by Targeting CCL7. Mol Ther 26: 582–592. PMC5835026

29. Grist, J. J., B. S. Marro, D. D. Skinner, A. R. Syage, C. Worne, D. J. Doty, R. S. Fujinami, and T. E. Lane. 2018. Induced CNS expression of CXCL1 augments neurologic disease in a murine model of multiple sclerosis via enhanced neutrophil recruitment. Eur J Immunol 48: 1199–1210. PMC6033633

30. Carter, S. L., M. Muller, P. M. Manders, and I. L. Campbell. 2007. Induction of the genes for Cxcl9 and Cxcl10 is dependent on IFN-gamma but shows differential cellular expression in experimental autoimmune encephalomyelitis and by astrocytes and microglia in vitro. Glia 55: 1728–1739.

31. Fitzgerald, D. C., B. Ciric, T. Touil, H. Harle, J. Grammatikopolou, J. Das Sarma, B. Gran, G. X. Zhang, and A. Rostami. 2007. Suppressive effect of IL-27 on encephalitogenic Th17 cells and the effector phase of experimental autoimmune encephalomyelitis. J Immunol 179: 3268–3275.

32. Lampron, A., A. Larochelle, N. Laflamme, P. Prefontaine, M. M. Plante, M. G. Sanchez, V. W. Yong, P. K. Stys, M. E. Tremblay, and S. Rivest. 2015. Inefficient clearance of myelin debris by microglia impairs remyelinating processes. J Exp Med 212: 481–495. PMC4387282

33. Das, A., M. Sinha, S. Datta, M. Abas, S. Chaffee, C. K. Sen, and S. Roy. 2015. Monocyte and macrophage plasticity in tissue repair and regeneration. Am J Pathol 185: 2596–2606. PMC4607753

34. Miron, V. E., A. Boyd, J. W. Zhao, T. J. Yuen, J. M. Ruckh, J. L. Shadrach, P. van Wijngaarden, A. J. Wagers, A. Williams, R. J. M. Franklin, and C. Ffrench-Constant. 2013. M2 microglia and macrophages drive oligodendrocyte differentiation during CNS remyelination. Nat Neurosci 16: 1211–1218. PMC3977045

35. Lloyd, A. F., and V. E. Miron. 2019. The pro-remyelination properties of microglia in the central nervous system. Nat Rev Neurol 15: 447–458.

36. Goldmann, T., and M. Prinz. 2013. Role of microglia in CNS autoimmunity. Clin Dev Immunol 2013: 208093. PMC3694374

37. Tay, T. L., D. Mai, J. Dautzenberg, F. Fernandez-Klett, G. Lin, Sagar, M. Datta, A. Drougard, T. Stempfl, A. Ardura-Fabregat, O. Staszewski, A. Margineanu, A. Sporbert, L. M. Steinmetz, J. A. Pospisilik, S. Jung, J. Priller, D. Grun, O. Ronneberger, and M. Prinz. 2017. A new fate mapping system reveals context-dependent random or clonal expansion of microglia. Nat Neurosci 20: 793–803.

38. Prajeeth, C. K., K. Lohr, S. Floess, J. Zimmermann, R. Ulrich, V. Gudi, A. Beineke, W. Baumgartner, M. Muller, J. Huehn, and M. Stangel. 2014. Effector molecules released by Th1 but not Th17 cells drive an M1 response in microglia. Brain Behav Immun 37: 248–259.

39. Saadi, F., D. Chakravarty, S. Kumar, M. Kamble, B. Saha, K. S. Shindler, and J. Das Sarma. 2021. CD40L protects against mouse hepatitis virus-induced neuroinflammatory demyelination. PLoS Pathog 17: e1010059. PMC8699621

40. Ransohoff, R. M., T. A. Hamilton, M. Tani, M. H. Stoler, H. E. Shick, J. A. Major, M. L. Estes, D. M. Thomas, and V. K. Tuohy. 1993. Astrocyte expression of mRNA encoding cytokines IP-10 and JE/MCP-1 in experimental autoimmune encephalomyelitis. FASEB J 7: 592–600.

41. Brambilla, R., P. D. Morton, J. J. Ashbaugh, S. Karmally, K. L. Lambertsen, and J. R. Bethea. 2014. Astrocytes play a key role in EAE pathophysiology by orchestrating in the CNS the inflammatory response of resident and peripheral immune cells and by suppressing remyelination. Glia 62: 452–467.

42. Sofroniew, M. V. 2009. Molecular dissection of reactive astrogliosis and glial scar formation. Trends Neurosci 32: 638–647. PMC2787735

43. Anderson, M. A., Y. Ao, and M. V. Sofroniew. 2014. Heterogeneity of reactive astrocytes. Neurosci Lett 565: 23–29. PMC3984948

44. Dutta, R., A. M. Chomyk, A. Chang, M. V. Ribaudo, S. A. Deckard, M. K. Doud, D. D. Edberg, B. Bai, M. Li, S. E. Baranzini, R. J. Fox, S. M. Staugaitis, W. B. Macklin, and B. D. Trapp. 2013. Hippocampal demyelination and memory dysfunction are associated with increased levels of the neuronal microRNA miR-124 and reduced AMPA receptors. Ann Neurol 73: 637–645. PMC3679350

45. Viola, A., F. Munari, R. Sanchez-Rodriguez, T. Scolaro, and A. Castegna. 2019. The Metabolic Signature of Macrophage Responses. Front Immunol 10: 1462. PMC6618143

46. Wang, F., S. Zhang, I. Vuckovic, R. Jeon, A. Lerman, C. D. Folmes, P. P. Dzeja, and J. Herrmann. 2018. Glycolytic Stimulation Is Not a Requirement for M2 Macrophage Differentiation. Cell Metab 28: 463–475 e464. PMC6449248

47. Hu, X., R. K. Leak, Y. Shi, J. Suenaga, Y. Gao, P. Zheng, and J. Chen. 2015. Microglial and macrophage polarization-new prospects for brain repair. Nat Rev Neurol 11: 56–64. PMC4395497

48. Touil, T., D. Fitzgerald, G. X. Zhang, A. Rostami, and B. Gran. 2006. Cutting Edge: TLR3 stimulation suppresses experimental autoimmune encephalomyelitis by inducing endogenous IFN-beta. J Immunol 177: 7505–7509.

49. Ellestad, K. K., S. Tsutsui, F. Noorbakhsh, K. G. Warren, V. W. Yong, Q. J. Pittman, and C. Power. 2009. Early life exposure to lipopolysaccharide suppresses experimental autoimmune encephalomyelitis by promoting tolerogenic dendritic cells and regulatory T cells. J Immunol 183: 298–309.

50. Buenafe, A. C., and D. N. Bourdette. 2007. Lipopolysaccharide pretreatment modulates the disease course in experimental autoimmune encephalomyelitis. J Neuroimmunol 182: 32–40.

51. Marta, M., A. Andersson, M. Isaksson, O. Kampe, and A. Lobell. 2008. Unexpected regulatory roles of TLR4 and TLR9 in experimental autoimmune encephalomyelitis. Eur J Immunol 38: 565–575.

52. He, Z., M. Riva, P. Bjork, K. Sward, M. Morgelin, T. Leanderson, and F. Ivars. 2016. CD14 Is a Co-Receptor for TLR4 in the S100A9-Induced Pro-Inflammatory Response in Monocytes. PLoS One 11: e0156377. PMC4881898

53. Bjork, P., A. Bjork, T. Vogl, M. Stenstrom, D. Liberg, A. Olsson, J. Roth, F. Ivars, and T. Leanderson. 2009. Identification of human S100A9 as a novel target for treatment of autoimmune disease via binding to quinoline-3-carboxamides. PLoS Biol 7: e97. PMC2671563 which is developing quinolines for commercial purposes. FI has a research grant from Active Biotech.

54. Liu, C., Y. Li, J. Yu, L. Feng, S. Hou, Y. Liu, M. Guo, Y. Xie, J. Meng, H. Zhang, B. Xiao, and C. Ma. 2013. Targeting the shift from M1 to M2 macrophages in experimental autoimmune encephalomyelitis mice treated with fasudil. PLoS One 8: e54841. PMC3572131

55. Wu, D., D. E. Sanin, B. Everts, Q. Chen, J. Qiu, M. D. Buck, A. Patterson, A. M. Smith, C. H. Chang, Z. Liu, M. N. Artyomov, E. L. Pearce, M. Cella, and E. J. Pearce. 2016. Type 1 Interferons Induce Changes in Core Metabolism that Are Critical for Immune Function. Immunity 44: 1325–1336. PMC5695232

56. Sriram, U., L. F. Barcellos, P. Villoslada, J. Rio, S. E. Baranzini, S. Caillier, A. Stillman, S. L. Hauser, X. Montalban, and J. R. Oksenberg. 2003. Pharmacogenomic analysis of interferon receptor polymorphisms in multiple sclerosis. Genes Immun 4: 147–152.

