## Supplementary materials for "IFN-induced protein with tetratricopeptide repeats 2 (Ifit2) limits autoimmune inflammation by regulating myeloid cell activation and metabolic activity"

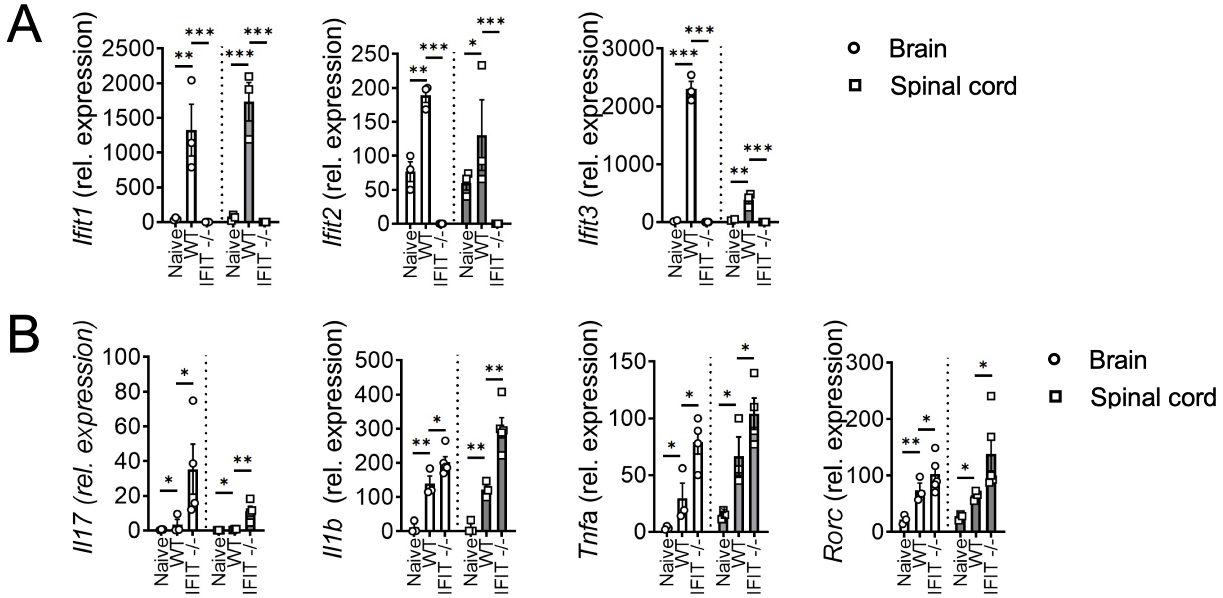


**Supplementary Fig 1.** ***Ifit*** **and cytokine mRNA expression in the brain and spinal cords of EAE-induced IFIT-WL KO mice.** Total RNA was extracted from brain and spinal cords tissue and reverse transcribed into cDNA. The expression of *Ifit1*, *Ifit2*, *Ifit3*, *Il17a*, *Il1b*, *Tnfa* and *Rorc* mRNA was determined by qPCR and normalized to *Gapdh.* n = 3-4 per group. *p < 0.05; **p < 0.01; ***p < 0.001; as determined by Mann-Whitney nonparametric test.

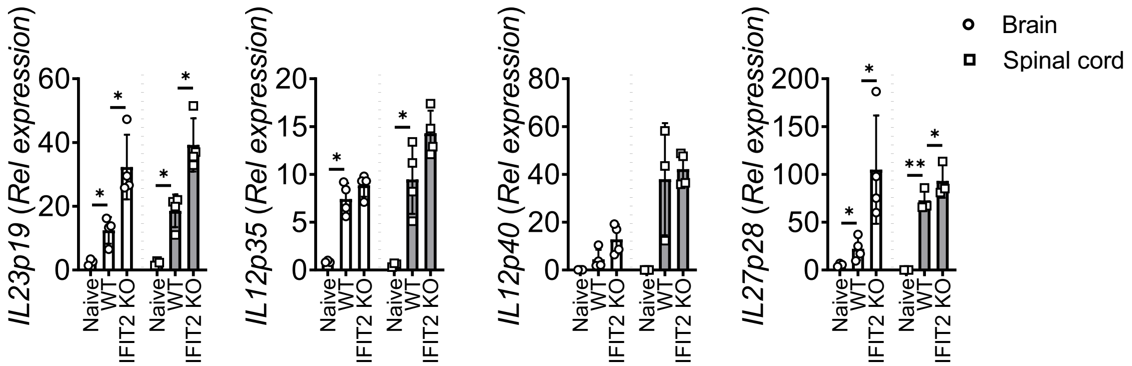


**Supplementary Fig 2. *IL-12* family cytokines expression in EAE-induced *Ifit2*$^{-/-}$ mice.** qPCR analysis of the indicated mRNAs in the brain and spinal cords from naïve, WT, and *Ifit2*$^{-/-}$ mice at the peak of disease (day 17 post immunization). Data were normalized by *Gapdh*. n = 3-4 per group. *$p < 0.05$; as determined by Mann-Whitney nonparametric test.

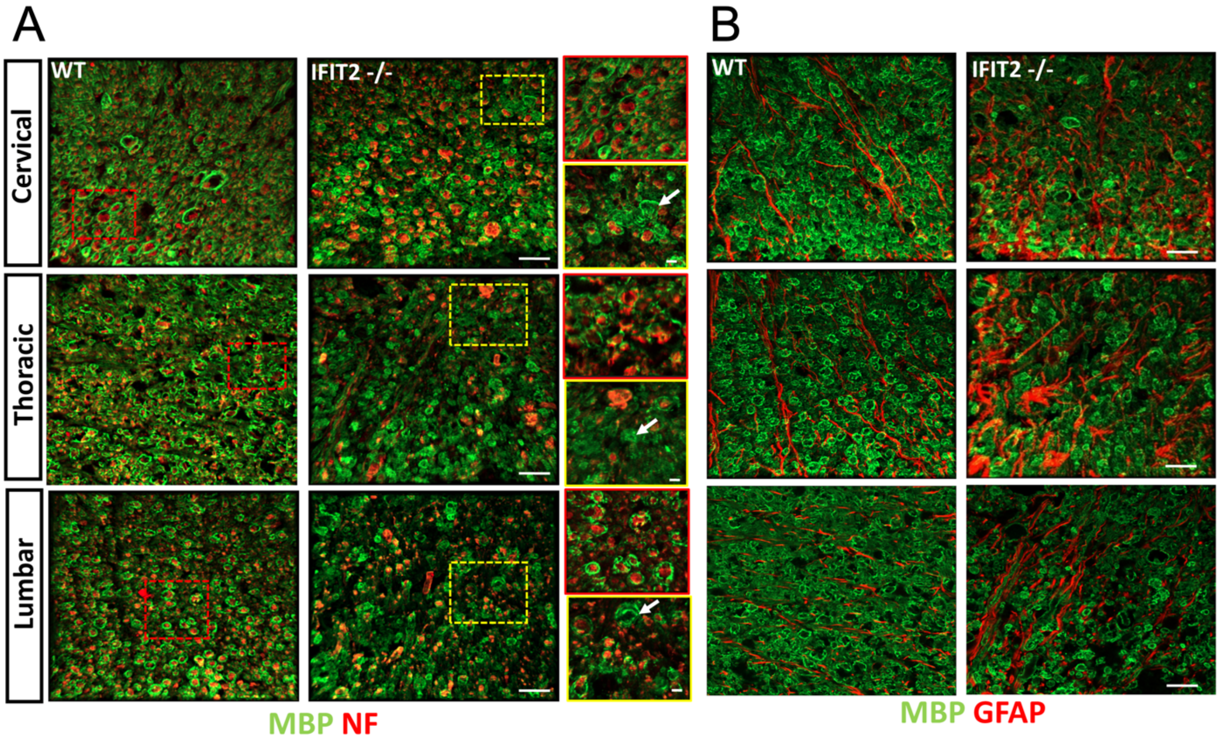


**Supplementary Fig 3. Ifit2 deficiency attenuates myelination, myelinated axon density but promotes astrogliosis in spinal cord of EAE mice**. (**A**) Myelination and myelinated axons were determined by staining for myelin basic protein (myelin, green) and neurofilament 200 kDa (NF200; axons, red). Representative 40X magnification images (Scale bar: 20µm) of ventral white matter showed decreased $MBP^+$ staining in *Ifit2*$^{-/-}$ mice. Enlarged images of inset showing decreased number of $NF200^+$ axons (red) surrounded by $MBP^+$ ring (arrow), indicating axonal degeneration. (B) Representative fluorescent images of GFAP-labeled astrocytes in spinal cord sections of WT and *Ifit2*$^{-/-}$ mice. There was notable increase in $GFAP^+$ cells in *Ifit2*$^{-/-}$ compared to WT at the peak of EAE.
